# Proteome analysis of xylose metabolism in *Rhodotorula toruloides* during lipid production

**DOI:** 10.1101/601930

**Authors:** Ievgeniia A. Tiukova, Jule Brandenburg, Johanna Blomqvist, Sabine Samples, Nils Mikkelsen, Morten Skaugen, Magnus Øverlie Arntzen, Jens Nielsen, Mats Sandgren, Eduard J. Kerkhoven

## Abstract

**Background:** *Rhodotorula toruloides* is a promising platform organism for production of lipids from lignocellulosic substrates. Little is known about the metabolic aspects of lipid production from the lignocellolosic sugar xylose by oleaginous yeasts in general and *R. toruloides* in particular. This study presents the first proteome analysis of the metabolism of *R. toruloides* during conversion of xylose to lipids.

**Results:** *R. toruloides* cultivated on either glucose or xylose was subjected to comparative analysis of its growth dynamics, lipid composition, fatty acid profiles and proteome. The maximum growth and sugar uptake rate of glucose-grown *R. toruloides* cells were almost twice that of xylose-grown cells. Cultivation on xylose medium resulted in a lower final biomass yield although final cellular lipid content was similar between glucose- and xylose-grown cells. Analysis of lipid classes revealed the presence of monoacylglycerol in the early exponential growth phase as well as a high proportion of free fatty acids. Carbon source-specific changes in lipid profiles were only observed at early exponential growth phase, where C18 fatty acids were more saturated in xylose-grown cells. Proteins involved in sugar transport, initial steps of xylose assimilation and NADPH regeneration were among the proteins whose levels increased the most in xylose-grown cells across all time points. The levels of enzymes involved in the mevalonate pathway, phospholipid biosynthesis and amino acids biosynthesis differed in response to carbon source. In addition, xylose-grown cells contained higher levels of enzymes involved in peroxisomal beta-oxidation and oxidative stress response compared to cells cultivated on glucose.

**Conclusions:** The results obtained in the present study suggest that sugar import is the limiting step during xylose conversion by *R. toruloides* into lipids. NADPH appeared to be regenerated primarily through pentose phosphate pathway although it may also involve malic enzyme as well as alcohol and aldehyde dehydrogenases. Increases in enzyme levels of both fatty acid biosynthesis and beta-oxidation in xylose-grown cells was predicted to result in a futile cycle. The results presented here are valuable for the development of lipid production processes employing *R. toruloides* on xylose-containing substrates.

## Background

Successful replacement of fossil diesel fuels with biological oils currently faces a number of challenges such as competing technologies, including electric vehicles, as well as unsustainable biological oils production practices such as clearing rainforests for palm oil cultivation (1, 2). There is also an increasing competition for lipids from the food sector due to a growing population, increasing standards of human nutrition, and sustainability concerns. Lipids are required in animal feeds, for instance oils from wild pelagic fish are used to feed fish in aquaculture. So-called “single cell oils” (SCOs) are lipids derived from microorganisms, which represent a potentially more sustainable source of lipids to partially replace fish-derived oils (3). Some microorganisms can produce lipids from lignocellulose-based substrates. The five-carbon (C5) monosaccharide xylose is the second most abundant sugar in plant biomass and its utilization by microorganisms is regarded as a challenge to reach a sustainable valorization of plant biomass and commercialization of lignocellulosic fuels and chemicals (4). Some oleaginous yeasts can convert xylose to lipids up to more than 20% of its dry cell weight: 22% *Rhodotorula glutinis* (5), 58% *Trichosporon fermentans* (6), 37% *Cryptococcus curvatus* (7) 33% *Cryptococcus albidus* (8), 42% *Rhodotorula toruloides* (9–11). Cultivation on xylose in combination with glucose was also investigated using *Lipomyces starkeyi* with 61% lipids of dry cell weight (DCW) produced (12), while engineering of *Yarrowia lipolytica* strain enabled it to assimilate xylose (13, 14). Microbial production of lipids from xylose was furthermore investigated in filamentous fungi: *Cunninghamella echinulata* (lipids constituted 58% DCW), *Mortierella isabellina* (lipids constituted 66% DCW) (15).

Establishment of sustainable lucrative microbial lipid production in industrial scale includes: i) use of complex substrate (generated from plant biomass and containing a mixture of C6 and C5- sugars and lignocellulose-derived inhibitors); ii) energy-consuming aerated cultivation; iii) production of oil with high value compositional fatty acid (FA) profile including n-3 long chain polyunsaturated FA; iv) production of value-added byproducts. The basidiomycete yeast *R. toruloides* is relatively fast growing, can assimilate pentoses (11), is resistant to lignocellulosic inhibitors (16), has a beneficial FA composition (9) including n-3 linolenic acid, and produces carotenoids (17). *R. toruloides* is one of the most promising species for lipid production with reported lipid yields of up to 65.5% DCW under high cell density cultivation with low-nitrogen feeding (18).

*R. toruloides* was originally applied in 1980 as a production organism for industrial-scale production of oil substitutes to replace cocoa butter (19). The mono-unsaturated 18-carbon FA oleic acid (C18:1 n-9) is the major FA and is accumulated to ~55% of total FA in *R. toruloides* Y4. Less than half of this amount, ~20 % composes palmitic acid (C16:0, saturated). Stearic (C18:0) and linoleic (C18:2 n-6) acid can each reach up to 10% of total FA (11, 18, 20). A number of *R. toruloides* strains have been subjected to genome sequencing (21–25), genetic engineering protocols are established (26–30) and the yeast is recognized as a novel platform organism for production of oleochemicals (31, 32). A number of cultivation conditions of *R. toruloides* have therefore been examined, e.g. growth on mixed substrates (33) or different cultivation modes on lignocellulosic substrates (34, 35) and lipid production at sulfate (36) or phosphate-limitation (37). Improved genetically modified strains of *R. toruloides* for conversion of biomass have been generated (38).

The molecular physiology of *R. toruloides* during lipid production from different sugar substrates have been investigated using both proteomics (39, 40) and transcriptomics (25, 41, 42) approaches. Meanwhile, the molecular physiology during conversion of xylose into lipids in *R. toruloides* and in oleaginous yeasts in general has been poorly investigated. Xylose assimilation involves the enzymes NAD(P)H-dependent xylose reductase (XR) and NADH-dependent xylitol dehydrogenase (XDH). XR reduces xylose to xylitol, which is then oxidized by XDH to xylulose. Xylulose is subsequently phosphorylated by xylulose kinase before entering the pentose phosphate pathway (PPP). *R. toruloides* also possesses the enzyme phosphoketolase, which cleaves xylulose-5-phosphate to acetyl phosphate and glyceraldehyde-3-phosphate (43). Unlike glucose assimilation, xylose assimilation requires NADPH due to the initial reduction of xylose into xylitol by XR. The cellular NADPH pool can be replenished by a number of pathways including PPP, the malic enzyme (ME) transhydrogenase cycle or the cytoplasmatic NADP^+^-dependent isocitrate dehydrogenase. While xylose utilization by oleaginous yeasts is poorly understood, a number of studies have investigated xylose metabolism in ethanol-producing yeasts: e.g. in engineered *Saccharomyces cerevisiae* (44), *Schefferomyces stipitis* (45), and *Ogataea polymorpha* (46). These studies reported that the carbon source affected levels of proteins involved in e.g. glucose repression, sugar transport, gluconeogenesis and amino acid catabolism. In the current study, comparative whole-proteome analysis was employed to study lipid formation in *R. toruloides* cultivated on either glucose or xylose as sole carbon source under conditions of nitrogen limitation.

## Results

### Batch cultivation of *Rhodotorula toruloides*

Growth, sugar consumption and lipid production of *R. toruloides* in medium containing either glucose or xylose as a carbon source were monitored in batch cultivations (Fig. 1). The C/N ratio was set to 75 in order to provide sufficient carbon surplus while averting inhibitory effect of nitrogen limitation. Sugar concentration was set at 40 g/l, similar to levels obtained in lignocellulose hydrolysate (47). Fermentations were performed at 25 °C to minimize energy input, which is beneficial for industrial processes (48). Cell growth was monitored by optical density (OD) and DCW measurements every eight hours. Samples for lipid and protein analysis were taken at early exponential phase (8 or 16 h), late exponential (16 or 40 hours) and lipid production phase (64 or 96 hours), with the later sampling-points referring to xylose, as indicated in Fig. 1.

**Fig. 1.**
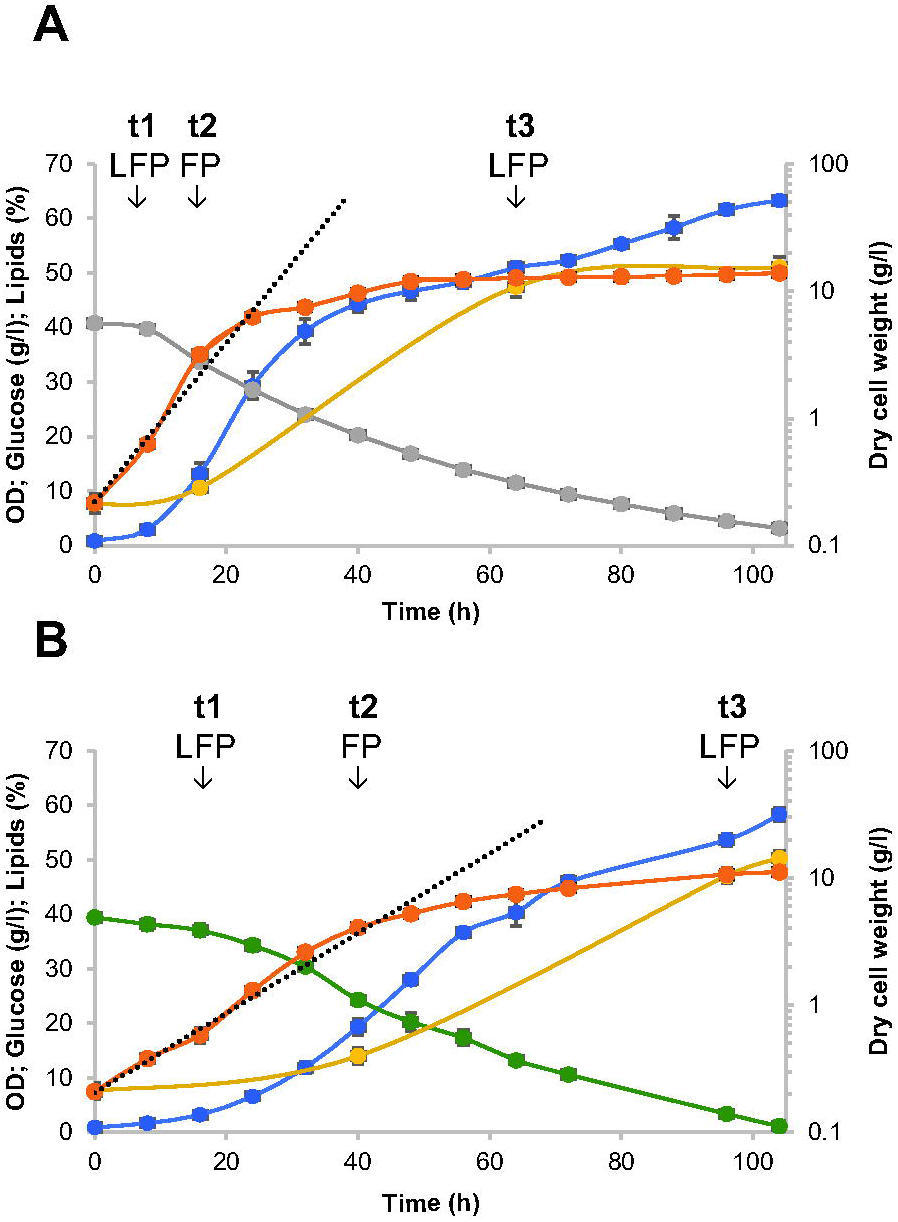
Batch cultivation of *R. toruloides* in chemically defined medium containing either glucose (A) or xylose (B). Cultivations of *R. toruloides* in a total volume of 0.5 1 medium were performed in triplicate at pH 5.5 and 25 °C in 0.7-1 bioreactors. Figure shows OD (blue), DCW (orange), glucose (grey), xylose (green), lipids (yellow), DCW trendline (interrupted grey line). Sample points for fatty acids profile (F), lipid classes (L) and proteome analysis (P) at early exponential (t1), late exponential (t2) and lipid production (t3) phase are all indicated. Mean values of three independent cultivation experiments are shown with error bars indicating one standard deviation.

The maximum growth rate of *R. toruloides* on glucose was almost twice that of xylose, which is also reflected in an uptake rate of glucose during exponential phase that is roughly double the xylose uptake rate (Table 1). The exponential growth phase on glucose lasted approximately 24 h compared to 40 h for xylose, after which the observed increase in DCW and OD slowed down (Fig. 1). The xylose level was lower than the glucose level after 72 h, even though the growth rate on xylose was lower. Possibly, nitrogen was depleted in xylose-based medium later than in the medium containing glucose (49–51). Growth on xylose resulted in lower final biomass yield and concentration as compared to glucose (Table 1, Fig. 1). This could indicate a higher ATP maintenance requirement of xylose-grown cells during the longer growth phase as compared to growth on glucose.

**Table 1.** Physiological parameters for batch cultivation of *R. toruloides* CBS 14 in minimal medium containing either glucose or xylose.

|  | Glucose | Xylose |
| --- | --- | --- |
| Lag phase (h) | 8 | 16 |
| Exponential phase (h) | 54 | 80 |
| $\mu_{\max}$ (h<sup>-1</sup>) | 0.14 | 0.07 |
| Final DCW (g/l) | 13.8 ± 0.13 | 11.1 ± 0.1 |
| Starting sugar (g/l) | 40.8 ± 0.3 | 39.4 ± 0.25 |
| Residual sugar (g/l) | 3.19 ± 0.81 | 1.1 ± 1 |
| Final $Y_{X/S}$ (gDW/g glucose<sup>-1</sup>) | 0.36 ± 0.004 | 0.28 ± 0.003 |
| Lipid content at t=0 (%) * | 7.59 ± 1.54 |  |
| Lipid content at t=2 (%) * | 10.58 ± 1.00 | 13.99 ± 1.5 |
| Lipid content at t=3 (%) * | 47.51 ± 1.81 | 47.05 ± 1.4 |
| Final $Y_{L/S}$ (g/g) | 0.19±0.01 | 0.15 ± 0.009 |

\* T=0 represents the beginning of cultivation, t=2 - late exponential growth phase, t=3 - lipid accumulation phase.

Significant lipid accumulation was observed at the end of the exponential growth phase, which was attributed to nitrogen depletion from the culture medium as commonly observed for cultivations of oleaginous yeasts (19). Although sugar levels decreased further after the late-exponential phase, particularly in glucose (Fig. 1, cf. glucose at 64 and 96 h), this only resulted in a slight increase in lipid content. The final lipid content under both growth conditions was approximately 51%, while final lipid yields were slightly higher on glucose, following the same trend as the biomass yield (Table 1). Interestingly, the higher lipid yield on glucose is comparable to yields reported previously (11), although the previous study reported similar biomass yields between glucose and xylose, which was not observed in the present study. In addition, the previous study reported lower lipid content of xylose-grown cells compared to glucose-grown cells. This disparity between studies could be explained by differences in the C/N ratio used for cultivation (cf. 65 (11) and 75 (our study)) or elemental medium composition (52). Meanwhile, *R. toruloides* CCT 0783 showed similar effects of glucose and xylose on the biomass yield, while lipid yields remained unaffected (34, 53).

### Lipid classes

Samples were taken for lipid class profiling to investigate whether the distribution of lipid classes changed during the cultivation (Fig. 1, Table 2). Monoacylglycerols (MAGs) were detected in the early exponential phase. MAGs are not commonly reported in yeast lipidomes (20, 54, 55), although it has been observed in cultivations of *R. toruloides* NP11 (56) and *Y. lipolytica* (57). The presence of MAGs in *R. toruloides* could be connected with diacylglycerol lipase activity, which cleaves one FA chain from diacylglycerols (DAGs) (58), as *Rhodotorula spp.* are used in the food industry as sources of lipases (59). Furthermore, a high proportion of free FA was observed in *R. toruloides* compared to other species where free FA are commonly below 5 % of total lipids (60, 61). MAGs were only detected during early exponential phase of the *R. toruloides* cultivations, while percentage of DAGs and free FA increased towards late exponential phase. This indicates that the absence of MAGs in late stage is not necessarily due to absence of lipase activity in *R. toruloides*.

**Table 2.** Lipid class composition of *R. toruloides* presented as percentage of total identified lipids. Samples were collected during early exponential phase (t1) and lipid production phase (t3). Total lipids were not determined at t1, but instead the values reported here are the mean total lipid contents between the first two total lipid measurements (Fig. 1), assuming a linear increase over time. Absolute levels of lipid classes were determined from the mean percentages of lipid classes and mean value of total lipid content. Abbreviations: DCW: dry cell weight, TAG: Triacylglycerols, PL: Phospholipids

|  | t1 |  |  |  | t3 |  |  |  |
| --- | --- | --- | --- | --- | --- | --- | --- | --- |
|  | Glucose<br>% of lipid<br>classes | mg/gDCW | Xylose<br>% of lipid<br>classes | mg/gDCW | Glucose<br>% of lipid<br>classes | mg/gDCW | Xylose<br>% of lipid<br>classes | mg/gDCW |
| Phospholipids | 15.09 ± 0.34 | 13.71 | 16.27 ± 1.5 | 17.56 | 4.11 ± 0.34 | 19.53 | 3.9 ± 0.21 | 18.37 |
| Monoglycerols | 1.02 ± 0.33 | 0.93 | 1.35 ± 0.5 | 1.46 | nd | nd | nd | nd |
| Diglycerols | 8.35 ± 0.7 | 7.59 | 7.88 ± 0.26 | 8.50 | 4.9 ± 0.3 | 23.28 | 4.02 ± 0.36 | 18.93 |
| Sterols | 14.11 ± 0.34 | 12.82 | 14.61 ± 0.79 | 15.76 | 6.49 ± 0.3 | 30.83 | 5.67 ± 0.15 | 26.71 |
| Free fatty acids | 38.15 ± 0.9 | 34.66 | 40.77 ± 3.1 | 43.99 | 11.9 ± 0.3 | 56.54 | 10.80 ± 0.25 | 50.87 |
| Triacylglycerols | 23.82 ± 1.7 | 21.64 | 19.39 ± 4.5 | 20.92 | 72.64 ± 0.76 | 345.11 | 75.61 ± 0.63 | 356.12 |
| TAG/PL | 1.58 |  | 1.19 |  | 17.66 |  | 19.37 |  |
| Total lipid content |  | 9.085±1 |  | 10.79±1 |  | 47.51±1.8 |  | 47.1±1.4 |

Lipid accumulation in *R. toruloides* is mainly in the form of neutral lipids as triacylglycerols (TAGs), irrespective of the carbon source (Table 2). The percentage of sterols was also approximately two-fold higher by the time the cells entered stationary phase compared to early exponential phase. In contrast, phospholipid proportions remained essentially unchanged, which was reflected by a pronounced increase in TAGs/phospholipids ratio compared to *S. cerevisiae* (54) or *Y. lipolytica* (60). The high TAGs/phospholipid ratio in *R. toruloides* suggested that the accumulated neutral lipids are stored in giant lipid droplets rather than the cell membrane. The *R. toruloides* lipid droplet proteome was shown to drive a formation of giant lipid droplets in this organism (56). Even at the start of cultivation, the observed TAGs/phospholipid ratio in *R. toruloides* was higher than that what has previously been reported for *S. cerevisiae* (54, 62) and *Y. lipolytica* (60, 63, 64).

No major differences in the proportions of lipid classes in *R. toruloides* were observed between cultivations containing either xylose and glucose. The high similarity in final relative content of TAGs in cells cultivated in medium containing glucose and xylose fits well with absence of differences in final total lipid content in examined cells.

### Fatty acids profiles

Samples were taken at different *R. toruloides* growth phases for lipid analysis to determine whether any changes in lipid classes and content correlates with changes in the FA profile (Fig. 1, Table 3). The dominant FA under all conditions was oleic acid. The final FA compositions agreed somewhat with previous reports (11, 18). The content of linoleic acid and stearic acid has previously been reported to be roughly 13 % for each of the total FA content in *R. toruloides* strains CBS 14 and Y4 (11, 18). However, in the present study we found that the amount of linoleic acid was almost 1.7 times lower than stearic acid. This difference can be due to differences in media composition and cultivation conditions: *R. toruloides* cultivations were carried out at 25°C in the present study compared to 30°C in previous reports (11, 18). Higher proportions of polyunsaturated FA have previously been reported to accumulate in *R. glutinis* when this yeast was cultivated at 25°C as compared to 30°C (65). Small amounts of heptadecenoic (C17:1) and heptadecanoic (C17:0) acid could also be detected in some samples (Table 3). Production of heptadecenoic acid has previously been reported in the two closely related species *Rhodotorula babjevae* and *Rhodotorula glutinis* (66, 67).

**Table 3.** Fatty acid profiles of *R. toruloides* during early exponential phase and lipid accumulation phase following batch cultivation in medium containing either glucose or xylose (presented as percentage of total identified, average ± standard deviation). Samples were collected at the beginning of cultivation (t0) as well as during early exponential phase (t1), late exponential phase (t2) and lipid production phase (t3). Polyunsaturated fatty acids were counted for the number of unsaturations according to (18).

|  | t0 | t1 |  | t2 |  | t3 |  |
| --- | --- | --- | --- | --- | --- | --- | --- |
|  | Glc = Xyl | Xylose | Glucose | Xylose | Glucose | Xylose | Glucose |
| C14:0 | 0.98 ± 0.25 | 1.03 ± 0.26 | 0.91 ± 0.03 | 1.23 ± 0.09 | 1.25 ± 0.05 | 1.28 ± 0.04 | 1.1 ± 0.07 |
| C16:0 | 14.35 ± 1.9 | 17.91 ± 2.5 | 17.16 ± 0.01 | 26.63 ± 0.22 | 25.76 ± 0.24 | 26.73 ± 0.26 | 25.55 ± 0.3 |
| C16:1 | 1.39 ± 0.28 | 0.96 ± 0.1 | 0.51 ± 0.02 | 0.98 ± 0.05 | 0.53 ± 0.01 | 1.00 ± 0.04 | 0.57 ± 0.05 |
| C17:0 | | 0.76 ± 0.13 | | | | | |
| C17:1 | 0.75 ± 0.18 | | | | | | |
| C18:0 | 3.84 ± 0.44 | 5.55 ± 0.57 | 7.63 ± 0.5 | 8.11 ± 0.28 | 9.31 ± 0.19 | 7.73 ± 0.22 | 7.83 ± 0.56 |
| C18:1(n-9) | 66.66 ± 2.28 | 46.27 ± 3.05 | 36.99 ± 1.3 | 45.26 ± 0.5 | 47.23 ± 0.61 | 45.07 ± 0.7 | 47.25 ± 1.06 |
| C18:2(n-6) | 6.62 ± 0.33 | 20.36 ± 4.16 | 23.56 ± 0.71 | 13.51 ± 0.69 | 11.58 ± 0.5 | 13.83 ± 0.57 | 13.06 ± 0.68 |
| C18:3(n-3) | 3.38 ± 0.86 | 5.74 ± 1.9 | 11.25 ± 0.98 | 2.78 ± 0.15 | 2.94 ± 0.22 | 2.99 ± 0.19 | 3.01 ± 0.25 |
| C24:0 | 0.9 ± 0.01 | 1.15 ± 0.13 | 0.66 ± 0.05 | 0.73 ± 0.05 | 0.76 ± 0.02 | 0.67 ± 0.05 | 0.91 ± 0.13 |
| Avg length | 17.4891 | 17.5942 | 17.4104 | 17.3038 | 17.3546 | 17.3084 | 17.3586 |
| Avg desaturation | 0.9218 | 1.0517 | 1.1837 | 0.816 | 0.7974 | 0.827 | 0.8297 |

The second major FA detected was either palmitic or linoleic acid, and this was growth phase dependent even while the average chain length remained relatively unchanged. Some differences in the FA chain lengths were observed compared to previous reports of the Y4 strain of *R. toruloides* (18), although the dominance of oleic acid, the average chain length and saturation levels were in agreement with published data.

Carbon source-specific changes in lipid profiles of *R. toruloides* were observed only at early exponential phase, where C18 FA were more saturated in xylose compared to glucose. Wiebe *et al.* observed a higher desaturation of C18 FA and an increase in palmitic acid proportion in response to xylose in *R. toruloides* CBS 14 (11), which was not observed in this study. Combined, it shows that medium composition and cultivation conditions (especially temperature) more significantly affect FA composition than carbon source and growth phase.

### Effect of growth phases on proteome composition of *R. toruloides*

As glucose and xylose assimilation proceed through distinct catabolic pathways, we anticipated that alternative carbon sources result in adjustments of the *R. toruloides* proteome. To investigate this, we examined relative proteome differences at different growth phases (Fig. 1). A total of 2995 proteins (corresponding to 36.7 % of the genome) could consistently be identified in at least two replicates of each tested condition (Additional file 2). Early and late exponential and stationary (lipid production) phases were compared separately for glucose and xylose cultures, in order to evaluate the effect of growth phase.

Of the 2995 identified proteins, 795 and 902 were differentially expressed when comparing early exponential and with stationary phase for xylose and glucose cultures, respectively (Additional file 1: Fig. S1, Additional file 3, 4). A core of 251 upregulated and 168 downregulated proteins were found when comparing lipid accumulation to early exponential phase in cells grown on both glucose and xylose, whereas a large part of the proteins were regulated specifically by carbon substrate (Additional file 1: Fig. S2).

Gene set analysis (GSA) performed to identify systemic changes revealed that genes involved in ribosome biogenesis and function (Additional file 1: Fig. S1) were downregulated during the cultivation on both sugars. These observed changes are more likely associated with the growth phase rather than lipogenesis, as transcriptional downregulation of ribosomes is connected to inactivation of the target of rapamycin (TOR) signalling pathway upon nitrogen depletion (68). A similar response has been reported for the *R. toruloides* strain NP11 as well as for the oleaginous yeast *Y. lipolytica* (69).

Proteins showing the largest differential expression during lipid accumulation phase on both glucose and xylose were identified. Proteins increasing 5- to 8-fold in abundance included urea transporter, NCS2 allantoate transporter, major facilitator superfamily (MFS) transporter of unknown specificity, siderochrome-iron transporter, amine oxidase, perilipin-like protein and aspartic proteinase (Additional file 1: Fig. S1, Additional file 3, 4). The observed upregulation of proteins involved in the assimilation of nitrogen agreed well with the expected cessation of the nitrogen catabolite repression (NCR) under conditions of nitrogen limitation (25). Regarding genes that are downregulated at the later growth phase, little consensus could be found between the two carbon sources (Additional file 1: Fig. S1, Additional file 3, 4).

### Carbon source dependent proteome changes

Comparison between carbon source for each time point resulted in the identification of 457, 230 and 181 proteins for early exponential, late exponential and lipid accumulation stage respectively (Additional file 1: Fig. S3, Additional file 5, 6, 7). It was observed that 43 upregulated and 6 downregulated proteins were common between all time points in xylose as compared to glucose grown cells (Additional file 1: Fig. S4). Additionally, 104 proteins were regulated similarly in response to xylose at early and late exponential phase, 5 proteins at late exponential and lipid accumulation phase and 7 proteins at early exponential and lipid accumulation phase. A majority of differentially expressed proteins were growth phase-specific between xylose and glucose.

Only limited conclusions could be drawn from gene-set analysis of these results (Fig. 2): proteins involved in ribosomal function were among the top upregulated proteins in early exponential phase on glucose, which is consistent with the observed higher growth rate on this carbon sources (Fig. 2, Table 1).

**Fig. 2.**
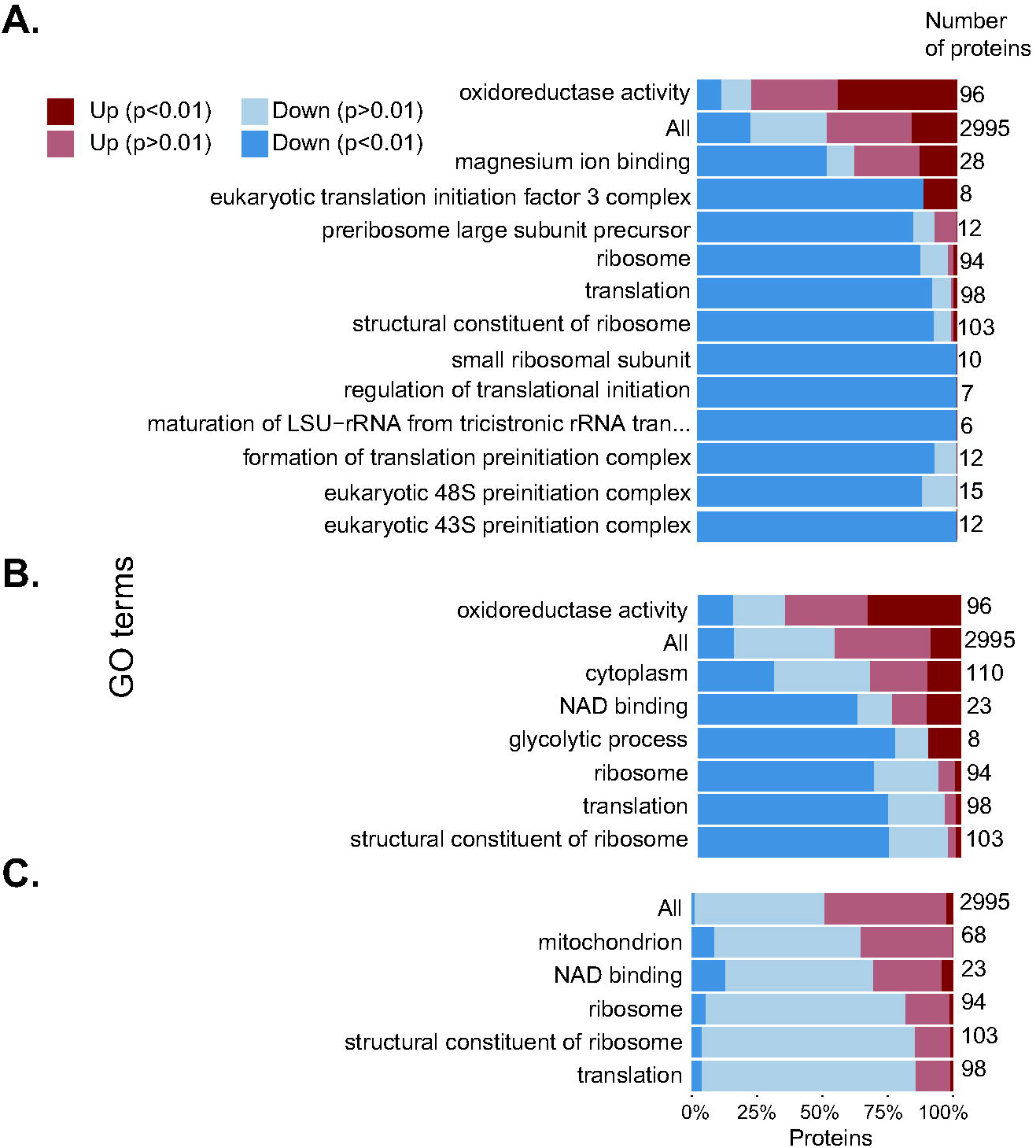
GO term enrichment analysis of differentially expressed proteins in *R. toruloides* grown on glucose or xylose during early exponential growth phase (A), late exponential growth phase (B), lipid accumulation phase (C). For each GO term showing significant change (rank score of ≥ 5), the direction and significance of the relative changes in protein levels are shown, together with the total number of proteins within each GO term.

Proteins involved in transport (MFS transporters) and metabolism of C5 sugar (xylitol dehydrogenase, L-iditol 2-dehydrogenase), NADPH generation (NADPH-dependent alcohol dehydrogenase; were among the top upregulated proteins on xylose at all time points (3- to 11-fold increase), while there were little similarities among the top downregulated proteins. Mitochondrial ME was among the top downregulated proteins in xylose grown yeast at early stationary phase.

As the untargeted GSA did not elucidate systemic transcriptional responses, we proceeded with targeted analysis of the largest observed transcriptional changes complemented with metabolic pathways and genes suspected to be affected by different carbon sources.

#### Sugar transport

The expression levels of various sugar transporters show a carbon source specific regulation. Thirteen putative sugar transporters were identified in the proteome dataset, of which eight (Rhto_01630, Rhto_03448, Rhto_07444, Rhto_06801, Rhto_06080, Rhto_01923, Rhto_00228, Rhto_07706) had significantly elevated levels (5.6- to 338-fold) for xylose cultivations as compared to glucose, suggesting their involvement in xylose transport (Fig. 3 and Additional file 8). Conversely, three transporters (Rhto_04266, Rhto_00984, Rhto_06016) with 16-fold elevated levels in cells fermenting glucose are likely involved in glucose transport. The relatively high level of the transporter Rhto_06080 irrespective of carbon source during exponential phase and Rhto_06016 and Rhto_01923 at stationary phase might indicate that these proteins function as unspecific sugar transporters. The transporter Rhto_01923 might have a high affinity to both sugars, while Rhto_06016 to xylose only. Lower levels of sugar transporters induced by glucose rather than xylose suggest high efficiency of expressed transporters.

**Fig. 3.**
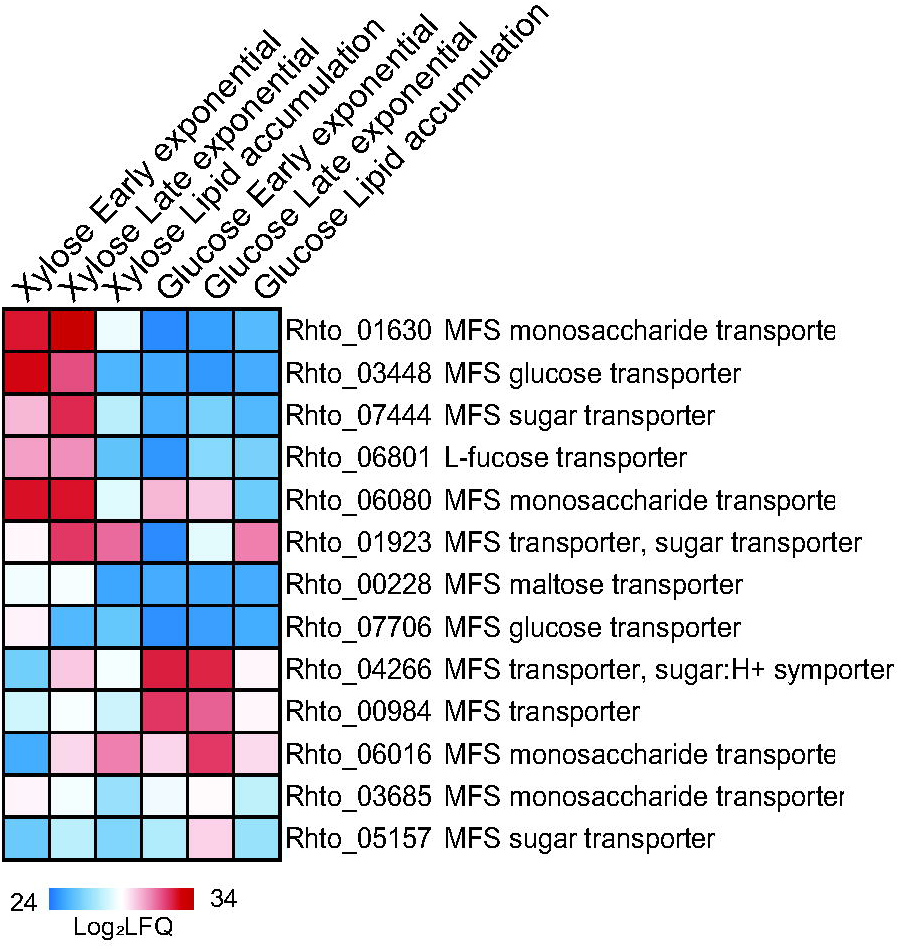
Heatmap showing relative abundance levels (log-transformed value of label-free quantification) of proteins involved in sugar transport during batch cultivation of *R. toruloides* in minimal medium containing either glucose or xylose.

#### Xylose assimilation

Proteins predicted to be involved in xylose assimilation were highly upregulated in the presence of xylose, in particular xylitol dehydrogenase (Fig. 4 and Additional file 8). Four NADPH-dependent XRs where strongly upregulated (6.8-68.5 fold) at all three time points during xylose cultivations. Besides xylose reductase Rhto_03963, also aldo/keto reductases Rhto_00641, Rhto_06555, Rhto_00602 are therefore likely involved in xylose assimilation.

**Fig. 4.**
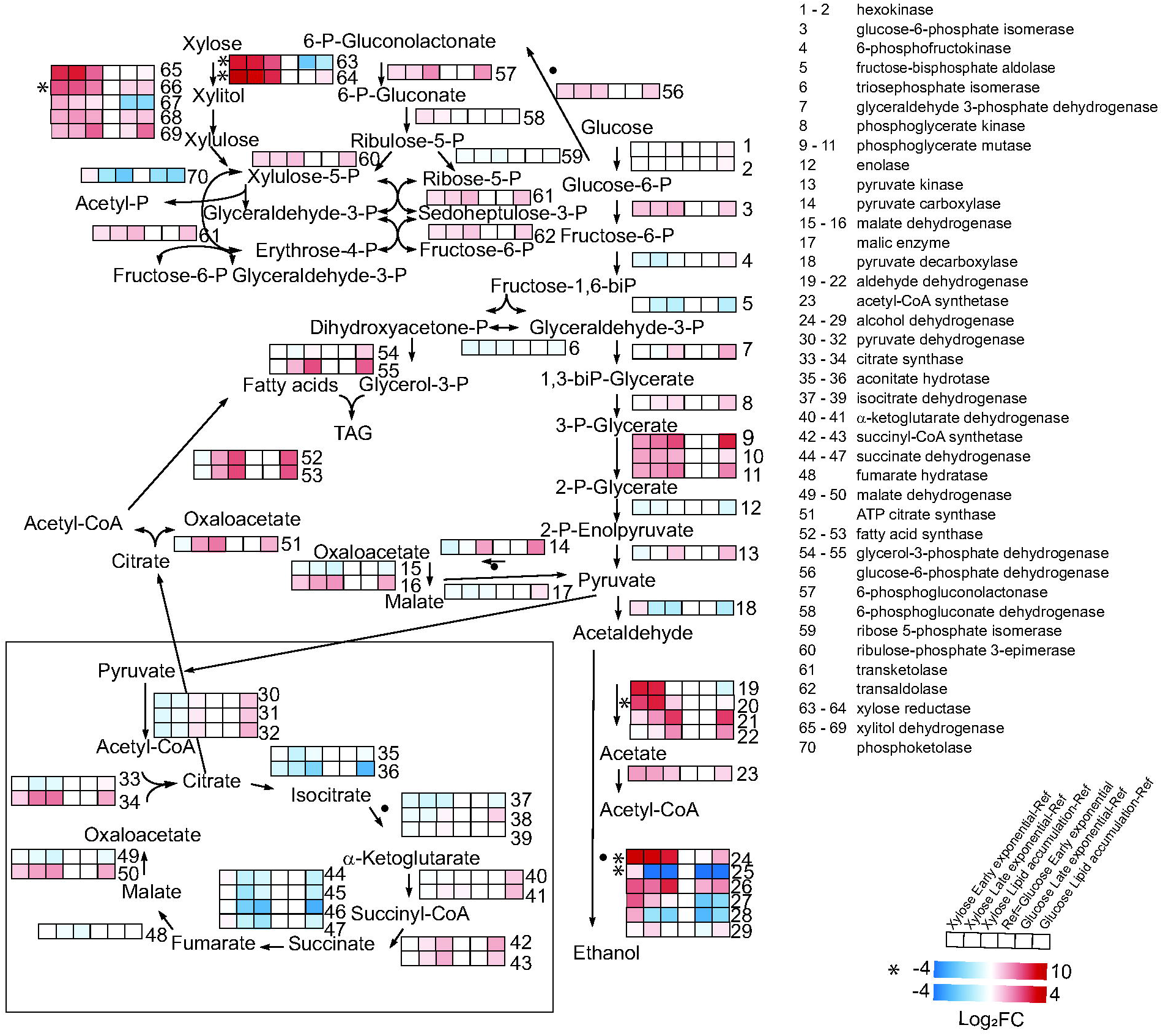
The figure shows a map of the central carbon metabolism of *R. toruloides* augmented with log_2_-transformed fold change of protein levels relative to the level at glucose early exponential phase. For most proteins, color scale ranges from −4 (blue) to +4 (red). For proteins with very large changes (*), the scale ranges from −4 (blue) to + 10 (red). More gene and protein names, annotations and expression abundance data are included in Additional file 8.

Apart from induction of proteins involved in xylose assimilation, the absence of glucose from the xylose-containing medium also resulted in a strong upregulation (64-1000 fold) of enzymes involved in assimilation of alternative carbon sources (beta-glucosidase Rhto_05582, mandelate racemase/muconate lactonizing enzyme Rhto_05818) (Additional file 5, 6, 7 and Fig. 4).

#### Glycolytic proteins and TCA cycle

Differences in expression levels of glycolytic and tricarboxylic acid (TCA) cycle proteins were less pronounced than those of sugar transporters, although this might be expected due to dominance of post-translational regulation of glycolytic enzymes in yeast (70, 71). Apart from upregulation of phosphoglycerate mutase during growth on xylose, no significant differences were observed in the levels of enzymes involved in glycolysis, PPP and phosphoketolase metabolic routes (Additional file 8, Fig. 4). From these data it was therefore not possible to infer whether xylulose-5-phosphate is metabolized via phosphoketolase or the non-oxidative steps of PPP and further glycolysis.

Oleaginous yeasts under nitrogen limitation are known to transfer citrate from the mitochondrion into the cytoplasm thereby reducing the flow of metabolites through the TCA cycle. In contrast to previous work (25) where higher C/N ratios were used, only aconitate hydrotase and succinate dehydrogenase showed moderate decreased levels during lipid accumulation in the present study.

#### NADPH generation

Each addition of a two-carbon unit during fatty acid elongation requires one molar equivalent of NADPH. The observed upregulation of PPP enzymes during lipid accumulation on both sugar substrates would suggest that NADPH generation in *R. toruloides* occurs mainly through PPP (Fig. 4 and Additional file 8). Such a scenario was further supported by the lack of significant differences in the levels of NAD(P)-dependent isocitrate dehydrogenase or cytosolic ME. Meanwhile, the upregulation of two other enzymes of the transhydrogenation cycle, i.e. pyruvate carboxylase and L-malate dehydrogenase (2.1-fold), suggests that this path has some contribution into regeneration of NADPH during lipogenesis.

In addition, we identified five aldehyde dehydrogenases (Rhto_05838, Rhto_04425, Rhto_04310, Rhto_06724, Rhto_04543) that might use NAD(P) as cofactor. A number of these dehydrogenases increased their expression during lipid accumulation (Additional file 8), while also some response to carbon source could be observed. Protein *Rhto_01922* (potentially mitochondrial), annotated as alcohol dehydrogenase (NADPH), was among the top upregulated proteins (832 times) on xylose at all time points.

#### Oleaginicity

The levels of ATP-citrate lyase (Rhto_03915), acetyl-CoA carboxylase (Rhto_02004) as well as both subunits of FA synthase (FAS) (Rhto_02139 and Rhto_02032) were upregulated in *R. toruloides* during conversion of both xylose and glucose into lipids (Additional file 8). Apart from cytoplasmic ME our results agree with earlier studies showing upregulation of key enzymes of lipid production machinery during lipogenesis (25, 39).

Levels of pyruvate decarboxylase decreased 2-fold during lipid accumulation on both carbon sources while levels of alcohol dehydrogenase (ADH; Rhto_03062) decreased sharply by 588 and 1024-fold on glucose and xylose, respectively. This suggests rechannelling of carbon towards more energy efficient carbon utilization, the pyruvate dehydrogenase—citrate synthase—ATP-citrate lyase path rather than to pyruvate decarboxylase—acetaldehyde dehydrogenase—acetyl CoA synthetase path.

#### Beta-oxidation

The *R. toruloides* proteomic data suggested that peroxisomal beta-oxidation triggered by nitrogen starvation was more pronounced in cells cultivated on xylose-based substrate than on glucose. The levels of enzymes involved in peroxisomal oxidation (Rhto_03890, Rhto_03776, Rhto_05407, Rhto_07118, Rhto_00300, Rhto_06581) was 8- to 16-fold higher in xylose than in glucose-grown cells already during the exponential growth phase (Additional file 8). The lipid production phase coincided with a decrease in levels of some enzymes of peroxisomal oxidation (Rhto_00300 and Rhto_04298) in xylose-grown cells while levels of others (Rhto_02517, Rhto_05520) increased in glucose-grown cells. These differences might indicate that TOR signaling is affected by carbon source in line with (72).

Similar trends were observed for mitochondrial beta oxidation: 8-128 times upregulation of the enzymes (Rhto_04971, Rhto_06738, Rhto_01625, Rhto_00397) happened in xylose grown cells already in the beginning of cultivation, however in glucose grown cells 2.3-8 times upregulation (Rhto_04971, Rhto_05797 and Rhto_00397) occurred only at lipid accumulation phase.

Simultaneous upregulation of the two inverse pathways beta-oxidation and FA biosynthesis in cells grown on xylose during induction of lipogenesis is expected to result in a futile cycle, which may partially explain the lower final biomass yields on xylose as compared to glucose.

#### TAG synthesis

The majority of enzymes involved in TAG synthesis did not change in protein levels during *R. toruloides* cultivation with the exception of a 4.1-fold increase in the level of glycerol-3-phosphate dehydrogenase (Rhto_02273) on both carbon sources as well as a 2-fold increase in the level of the phosphatidic acid biosynthetic 1-acyl-sn-glycerol-3-phosphate acyltransferase (Rhto_06718) in glucose-grown cells (Additional file 8). An increase in TAG lipase (Rhto_00361) level was observed during the lipid accumulation phase on xylose, that is probably counteracted by increase of perilipins, which protect lipids in droplets from degradation. The observed increase of TAG coincided with elevated absolute levels of DAGs and free FAs during stationary phase. However, MAGs were only detected during early exponential phase.

#### Phospholipids

Observed differences in levels of phospholipid biosynthetic enzymes suggest that phospholipid biosynthesis is affected by the choice of carbon source. Xylose-grown cells displayed a 2.5-fold increase in phosphatidate cytidylyltransferase (Rhto_01718) levels (converting phosphatidic acid to CDP-DAG), 2-fold upregulation of phosphatidyl-*N*-methylethanolamine *N*-methyltransferase Rhto_03783 (forming phosphatidyl-*N*-dimethylethanolamine) and 2.3-fold downregulation of CDP-diacylglycerol-inositol 3-phosphatidyltransferase (Rhto_02741, synthesising phosphatidyl-1D-myo-inositol) during lipid production phase (Additional file 8). In glucose-treated cells the level of phosphatidylserine decarboxylase Rhto_03399 forming phosphatidylethanolamine was upregulated 16 times. Meanwhile, analysis of lipid classes did not reveal any difference in proportion of phospholipids in *R. toruloides* cells in response to carbon source.

#### Mevalonate pathway

The mevalonate pathway is involved in the synthesis of dolichol, carotenoids and sterols from for acetyl-CoA and ATP and is therefore in competition with FA biosynthesis. Some enzymes of this pathway were regulated differentially in *R. toruloides* in response to carbon source. For instance, during the lipid production phase in glucose-treated cells, the level of acetyl-CoA C-acetyltransferase Rhto_02048 decreased 3.5-fold. Xylose-grown cells displayed a 2-fold increase in the enzyme geranylgeranyl diphosphate synthase (Rhto_02504, Additional file 8). Phytoene dehydrogenase (Rhto_04602) was upregulated in stationary phase cells irrespective of carbon source.

#### Amino acids biosynthesis

While we did not anticipate that carbon source would affect other metabolic pathways, we did observe changes in e.g. amino acid biosynthesis. A 2.5-fold decrease in the levels of leucine biosynthetic enzymes was observed in both xylose- and glycose grown cells upon the initiation of the lipid production phase (Additional file 8), while the downregulation of enzymes involved in the biosynthesis of proline (proline synthetase associated protein Rhto_04810) and leucine (branched-chain-amino-acid aminotransferase Rhto_05760 and Rhto_08045, acetolactate synthase Rhto_03298 and acetolactate synthase small subunit Rhto_03988) was detected in xylose-as compared to glucose-grown cells at the exponential phase. This might be related to the decrease of protein biosynthesis in xylose-treated cells.

The downregulation of enzymes involved in the synthesis of leucine (3-isopropylmalate dehydratase Rhto_07040, branched-chain-amino-acid aminotransferase Rhto_05760, acetolactate synthase Rhto_03298, ketol-acid reductoisomerase Rhto_04566) and the interconversion of methionine and cysteine (cystathionine beta-lyase Rhto_02122) was observed during lipid production in glucose-grown *R. toruloides* cells compared to earlier growth phase. Lipid accumulation in *Y. lipolytica* has previously been shown to be accompanied by downregulation of leucine biosynthesis (69).

#### Oxidative stress response

Levels of proteins involved in glutathione metabolism increased in *R. toruloides* cells cultivated on xylose but also during lipid production, possibly as a response to oxidative stress induced under these conditions. Levels of lactoylglutathione lyase (Rhto_06289), glutathione peroxidase (Rhto_00225), glutathione *S*-transferases (Rhto_03923, Rhto_00450) and NAD(P)H-dependent *S*-(hydroxymethyl) glutathione dehydrogenase (Rhto_03559) increased during exponential phase in xylose-grown cells compared to glucose-grown cells (Additional file 8). Levels of lactoylglutathione lyase (Rhto_06289), glutathione *S*-transferases (Rhto_00450, Rhto_03923) and glutathione peroxidase (Rhto_00225) also increased during the lipid production phase in xylose-grown cells relative to exponential growth phase. The levels of glutathione peroxidase (Rhto_00225) and glutathione *S*-transferase (Rhto_03923) also increased during the lipid accumulation phase in glucose-grown *R. toruloides* cells. Formate dehydrogenase (Rhto_06042) was detected among top upregulated enzymes during cultivation in xylose-based medium. Formate dehydrogenase is involved in the oxidation of formic acid to carbon dioxide as the final step of glutathione-dependent formaldehyde detoxification (73).

Activation of enzymes involved in carotenoid biosynthesis during lipid accumulation might be connected with oxidative stress response. Phytoene dehydrogenase Rhto_04602 was upregulated during lipid accumulation as compared to exponential growth phase in *R. toruloides* cells grown on both substrates (Additional file 8). Additionally, we detected increased levels of geranylgeranyl diphosphate synthase Rhto_02504 in glucose-grown cells.

#### Nitrogen starvation response

A number of parallel changes were observed in both glucose- and xylose-grown *R. toruloides* cells in response to nitrogen starvation. The quicker depletion of nitrogen from glucose-containing medium meant that differences in levels of proteins involved in the nitrogen starvation response were first observed in glucose-grown cells during exponential phase. Therefore the levels of amino acid transmembrane transporters (Rhto_01344, Rhto_00398, Rhto_07341) and proteins involved in amino acid degradation (branched-chain amino acid aminotransferase, Rhto_05760; tyrosine aminotransferase, Rhto_01136; aromatic amino acid aminotransferase I, Rhto_07034; L-asparaginase, Rhto_07901) appeared lower in xylose-grown *R. toruloides* cells compared to glucose-grown cells during early exponential phase as the xylose-grown cells had not yet fully exhausted their nitrogen source and hence not activated their nitrogen starvation response (Additional file 8). Further depletion of nitrogen from the medium during the cultivation in xylose eventually triggered a nitrogen starvation response that was consistent between the two carbon sources, such as increased levels of glutamate dehydrogenase, allantoicase, urea and amino acid transporters and autophagy related proteins. Nitrogen starvation induces autophagic processes, liberating intracellular space for growth of lipid droplet (25, 53). We observed upregulation of several autophagic proteins Rhto_05541, Rhto_01208, Rhto_07813 in *R. toruloides* cells at lipid accumulation as compared to exponential phase on both substrates (Additional file 8). Additionally, at this stage in xylose-grown cells the levels of autophagy-related protein Rhto_06526, Rhto_02366 and Rhto_07138 were upregulated. Interestingly, several proteins involved in autophagy Rhto_06526, Rhto_01636 and Rhto_05541 were upregulated in cells cultivated in xylose as compared to glucose at early exponential phase. This might indicate stress response to the absence of glucose.

## Discussion

A number of previous proteomic and transcriptomic studies have investigated metabolism of xylose in ethanol-producing yeasts (44–46), while this study of *R. toruloides* represents the first proteomic investigation of an oleaginous yeast during conversion of xylose to lipids. As the major constituent of hemicellulose, xylose is the second most abundant monosaccharide in nature after glucose (4).

Many notable differences were observed between the proteomes of cells cultivated in medium containing either glucose or xylose. Glucose- and xylose-grown cells had similar lipid content and profiles, but xylose-grown cells displayed lower biomass yields, in agreement with previous observations (34). The growth rate of xylose-grown *R. toruloides* cells in exponential phase was roughly half of the growth rate on glucose. The observed decrease in the levels of ribosomal proteins and other proteins associated with translation of mRNA in xylose-grown cells was the most prominent change in proteome between the two sugar substrates and is likely rather connected with differences in the growth rates.

The lower biomass yield and growth rate are possibly an effect of higher ATP demands by *R. toruloides* when grown on xylose. In the presence of excess sugar, and in particular during lipid accumulation, beta-oxidation is a futile cycle, and as proteins involved in beta-oxidation were upregulated during growth on xylose it is feasible that this futile cycle is underlying the higher ATP requirements. This is potentially an effect of absence of glucose, which would otherwise down-regulate beta-oxidation. Abolition of this pathway is therefore expected to have a significant beneficial impact on not only lipid formation but also on growth in excess of sugar with more pronounced effect on xylose as compared to glucose. Disruption of peroxisome biogenesis in *Y. lipolytica* has previously been reported to impair β-oxidation, which resulted in increased lipid accumulation from glucose (74). However, another study showed that disruption of peroxisome biogenesis in *R. toruloides* had the opposite effect, namely a decreased lipid production in glucose medium (75).

There were indications of that sugar transport is the underlying reason for the observed differences in physiology between glucose- and xylose-grown *R. toruloides* cells. A number of candidate glucose- and xylose-specific transporters were identified from the *R. toruloides* proteome datasets. As assimilation on xylose required strongly increased levels of some transport proteins, while only a reduced growth rate could be obtained, it is suggested that the xylose transporters are less efficient than those responsible for glucose transport. It is possible that the applied protein extraction methods have a bias against membrane proteins; however, improving xylose transport could be a valuable strategy for further optimization of *R. toruloides* for use of lignocellulosic biomass.

We have detected expression of *S. cerevisiae* homologs of members of glucose catabolite repression pathway in *R. toruloides*. However, apart from the monosaccharide transporter, Rhto_06016 (homolog of *SNF3*), we did not find any homologs to *S. cerevisiae* proteins-members of glucose induction pathway in genome of *R. toruloides*, which in the absence of glucose represses proteins required for glucose assimilation, such as glucose transporters (76). Our results showed that expression of some of glucose transporters was lower in xylose-grown cells. It remains unclear which mechanisms in *R. toruloides* provide function of glucose induction pathway. Possibly, release from strong inhibition of glucose-induced proteins can provide high fitness of microorganism in nutrient poor environment, such as soil and phyllosphere - natural habitats of *R. toruloides* (77).

Analysis of metabolic routes that can be used by *R. toruloides* to assimilate sugars did not elucidate a dominant pathway. As the main catabolic flux of glucose is funneled through glycolysis, this results in a lower flux through xylulose-5-P compared to catabolism of xylose. From this reduced availability of xylulose-5-P one might expect a downregulation of phosphoketolase during growth on glucose, however, its protein level showed no significant change as response to the carbon source. One possibility is that once cells pre-cultured on glucose were transferred to xylose medium, the xylose flux into cells was low enough that PPP enzymes levels did not have to be increased further to ensure efficient xylose catabolism. The phosphoketolase reaction normally enables higher efficiency of carbon metabolism since bypasses the wasteful decarboxylation pyruvate to acetyl-CoA, which causes a loss of one third of the carbon substrate, while phosphoketolase can produce an acetyl residue directly from a C5-substrate. However, under conditions of excess of carbon and simultaneous nitrogen limitation, the carbon saving phosphoketolase reaction may become unnecessary, which could result in xylulose-5-P being redirected towards PPP. Such a scenario would also explain the decrease in phosphoketolase levels during the lipid production phase on both substrates since the NADPH-regenerating reactions of PPP occur upstream of xylulose-5-P and therefore phosphoketolase is not expected to be regulated by NADPH demand. It was speculated in a previous study that assimilation of xylose via phosphoketolase would enable slightly higher lipid yields when grown on xylose (0.34 g/g sugar) as compared to growth on glucose (0.32 g/g) (78). It should be noted that partial assimilation of xylose via PPP would still be required for synthesis of essential nucleic acid precursors as well as regeneration of NADPH for lipogenesis.

NADPH generation for lipid synthesis was assumed to occur through PPP although the observed increase of the levels of transhydrogenase pathway enzymes indicates that ME could contribute to NADP^+^ reduction during stationary phase. Previous studies showed that in other oleaginous yeasts, *Y. lipolytica* and *L. starkeyi* PPP provides major contribution to NADPH pool during lipogenesis (79, 80). It has been previously demonstrated, that overexpression of genes encoding native ME slightly increased lipid accumulation in *R. toruloides* (75). Interestingly, an increase in levels of a putative NADP^+^-dependent alcohol dehydrogenase was observed, which suggests that oxidation of one or more alcohols into the corresponding aldehydes could contribute to supply of NADPH pool. Aldehyde dehydrogenase could also potentially generate NADPH, but the substrate specificity of this enzyme must first be determined. Lipid synthesis is an NADPH demanding process, but the absence of significant changes in the levels of NADPH regenerating enzymes in the present study would suggest that NADPH availability is not limiting lipid accumulation in xylose-grown *R. toruloides* cells.

Additional metabolic processes displaying significant changes in protein levels between glucose- and xylose-grown *R. toruloides* cells included biosynthesis of amino acids and phospholipids as well as oxidative stress response. Similar effect of xylose on ethanol producing yeasts has been reported previously (44–46). Some of these effects, such as downregulation of leucine biosynthesis and upregulation of xylose transporters during growth on xylose were reminiscent of proteomic changes observed in ethanol producing yeasts during cultivation on either xylose or glucose. The increased levels of two of the three enzymes involved in the glutathione-dependent formaldehyde detoxification pathway during cultivation in xylose-based medium was curious, as there was no obvious source of formaldehyde formation equivalent to what occurs during assimilation of methylated substrates such as methanol or methylated amines.

As the difference in *R. toruloides* growth rate were likely underlying many of the detected protein levels, it would be beneficial to perform continuous cultivations on both carbon sources to provide additional insight on the response on protein levels. Future studies should characterize proteome of *R. toruloides* cells with higher lipid content generated from xylose-based substrate. In addition, multicomponent lignocellulose-based medium could provide valuable information for further optimization of hemicellulose fermentation by *R. toruloides*. While proteomics measurements are instrumental to inform how transcriptional regulation affects metabolism, further augmenting with transcriptomics analysis could provide a higher depth of analysis due to its larger coverage of the total genome.

## Conclusions

This study is the first investigation of proteome of oleaginous yeast *R. toruloides* during conversion of xylose into lipids. Our study showed two times lower maximum growth rate and sugar uptake rate of *R. toruloides* cells in xylose as compared to glucose-based substrate. Proteome analysis revealed significantly lower levels of ribosomal proteins and translation associated factors in xylose-grown cells as compared to glucose-grown cells. In addition, xylose-grown cells contained higher levels of enzymes involved in peroxisomal beta-oxidation and oxidative stress response and lower levels of enzymes involved in leucine biosynthesis compared to cells cultivated on glucose. The levels of enzymes involved in sugar transport and phospholipid biosynthesis differed in response to carbon source. In addition, the proteomic data reported here identifies several potential target sugar transporters (Rhto_01630, Rhto_03448, Rhto_07444, Rhto_06801, Rhto_06080, Rhto_01923) and enzymes involved in peroxisomal beta-oxidation (Rhto_03890, Rhto_03776, Rhto_05407, Rhto_07118, Rhto_00300, Rhto_06581) for genetic engineering in order to optimize xylose metabolism.

## Abbreviations

PPP: pentose phosphate pathway
SCOs: single cell oils
DCW: dry cell weight
FA: fatty acid
XR: xylose reductase
XDH: xylitol dehydrogenase
ME: malic enzyme
OD: optical density
MAGs: monoacylglycerols
DAGs: diacylglycerols
TAGs: triacylglycerol
GSA: gene set analysis
TOR: target of rapamycin
FAS: fatty acid synthase
ADH: alcohol dehydrogenase
MFS: major facilitator superfamily
NCR: nitrogen catabolite repression
TCA: tricarboxylic acid
HPLC: high performance liquid chromatography
GC: gas chromatography
TLC: thin-layer chromatography
LC-MS: liquid chromatography-mass spectrometry

## Methods

### Strain and Media

*R. toruloides* CBS 14 was obtained from the Westerdijk Fungal Biodiversity Institute (Utrecht, the Netherlands) and stored at −80°C in 50% v/v glycerol. For cultivation experiments, frozen *R. toruloides* stocks were revived on solid YPD medium (20 g/l glucose, 20 g/l peptone and 10 g/l yeast extract, 16 g/l agar) and incubated for 3 days at 25°C. *R. toruloides* was precultured in 200 ml YPD liquid medium in 500 ml Erlenmeyer flasks and incubated for 3 days at 25°C on an orbital shaker at 150 rpm. The chemically defined cultivation medium used for growth experiments contained: 1.173 g/l (NH_4_)_2_SO_4_; 1.7 g/l yeast nitrogen base without amino acids or ammonium sulfate; 3 g/l KH_2_PO_4_; 0.5 g/l MgSO_4_-7H_2_O; 40 g/l carbon source (either xylose or glucose). The carbon/nitrogen (C/N) ratio of the medium was set to 75. 0.07% (v/v) polypropylene glycol was used to prevent adhesion and foaming of yeast cells to glass surfaces within the bioreactors.

### Yeast cultivation

Batch cultivations were performed in 0.7-l bioreactors (Multifors 2, Infors-HTBottmingen, Switzerland). Cultivation experiments for each carbon source (glucose and xylose) for analysis of lipid accumulation were performed in triplicate. *R. toruloides* precultures were washed twice with a saline solution (9 g/l NaCl) and then used to inoculate the bioreactors to a final optical density at 600 nm (OD_600_) of approx. 1.0 in a total volume of 0.5 1 chemically defined cultivation medium containing either glucose or xylose as carbon source. Batch cultivations were performed at pH 5.5 and 25 °C. The dissolved oxygen tension (DOT) was kept above 20 by regulating a stirring speed between 200 and 600 rpm at an aeration rate of 1 l/min. Samples for high performance liquid chromatography (HPLC) analysis as well as OD_600_ and DCW measurements were taken every 8 hours. Samples for lipid analysis were taken at early exponential phase, late exponential phase and early stationary phase for each cultivation experiment.

### Analytical methods

Cell OD_600_ was measured using an Ultrospec 1100 pro spectrophotometer (GE Healthcare, Chicago IL). Yeast cell biomass was determined by cell dry weight measurement from 2 ml of culture samples. The samples were centrifuged at 17,000 × *g*, washed twice with deionized water, incubated in an oven at 105°C for 72 h and then weighed. Each DCW measurement was done in triplicate. HPLC measurements of ethanol, acetate and glycerol were performed as described previously (81). Yeast lipids were extracted using the method described by Folch et al. (82) with some modifications (83). Briefly, 100 mg of freeze-dried yeast cells were suspended in 1 ml 1 M HCl and soaked for 15 min followed by 1 h incubation at 75 °C. 5 ml of chloroform:methanol (2:1, v/v) was added to the sample tube, which was then vortexed for 15 min at maximum speed. The sample tube was centrifuged at 5,000 × *g* for 3 min at room temperature followed by the transfer of the lower lipid layer to a tube of known weight. The extraction was repeated with remaining upper layer of the sample. The lipid phases from the first and second extractions were pooled followed by evaporation of the solvent under N_2_ gas flow. The total amount of lipids was determined gravimetrically. Total lipid content was determined at the start of fermentation instead of at early exponential phase due to a lack of material.

Dried lipid samples were dissolved in 0.5 ml hexane and methylated with BF_3_ according to Appelqvist (84). Methyl esters were analysed by gas chromatography (GC) as described previously (83, 85). The standard mixture 68A (Nu-Check, Elysian, MN) and retention times were used to identify FA. The double binding or unsaturation index (UI) was calculated as UI [%] = [%16:1 + %17:1 + %18:1 + (%18:2)·2 + (%18:3)·3]/100.

Lipid class composition was determined according to Olsen & Henderson (1989) (86) with slight modifications. Samples were diluted to a final concentration of 1 g/l in hexane, and 5 µl of each sample was then applied with a CAMAG thin-layer chromatography (TLC) Sampler ATS4 (Camag, Switzerland) 2 cm from the base edge of the TLC plates (pre-coated with silica gel TLC plates (20 × 10 cm; Silicagel 60; 0.20 mm layer, Merck, Darmstadt, Germany) in 2 mm bands with an application speed of 250 nl/s. N_2_ was used as spray gas. All samples were applied in duplicate, and the distance between tracks was 9.8 mm. Separation of the lipid classes was executed with a CAMAG Automatic Developing Chamber 2 (ADC 2) (Camag, Switzerland). Lipid classes were separated using a hexane:diethyl ether:acetic acid (85:15:2; v/v/v) mobile phase. Afterwards, separation procedure plates were submerged in a solution of 3 % (w/v) cupric acetate in 8 % (v/v) phosphoric acid and then charred for 20 min at 140 °C. Quantitative analysis of the separated lipid classes was done by scanning the plates with a CAMAG TLC Scanner 3 (Camag, Switzerland). The scanning was performed at a speed of 20 mm/s and a data resolution of 100 μm/step, with a slit dimension of 6.00 × 0.45 mm at a wavelength of 350 nm. Lipid classes were identified by comparison to external standards (TLC 18-4A, Nu-Chek Prep, Elysian, USA; Ergosterol, PHR1512, Sigma-Aldrich, Sweden). The mode Savitsky-Golay 7 was used for data filtering. Manual baseline and peak correction were used when necessary.

### Preparation of samples for proteome analysis

Samples for proteome analysis were taken at early exponential phase (8 or 16 h), late exponential (16 or 40 hours) and lipid production phase (64 or 96 hours), with the later sampling-points referring to xylose. Cell pellets were washed and frozen. Cell pellets were thawed and resuspended in 500 μl of a solution of 4 % (w/v) SDS, 10 mM DTT and 100 mM Tris-HCl (pH 8). Resuspended cell pellets were then transferred to 2 ml FastPrep tubes containing ca 500 μl of a 3:1 (w/w) glass bead mixture of bead sizes 106 μm (Sigma Aldrich, product no. G4649) and 425-600 μm (Sigma Aldrich, product no. G8772), respectively. Cells were disrupted using a FastPrep homogenizer (MP Biomedicals, CA) with three 45-s cycles at full speed interspersed with 5 min cooling on ice between runs. The tubes were then centrifuged for 15 min at 20,000 × *g*, and the supernatant was transferred to fresh tubes. An upper red layer of the supernatant could be observed at this point and was carefully avoided. The bottom layer was used to generate tryptic peptides according to the Filter Aided Sample Prep (FASP) protocol (87). Briefly, 50 μl of each FastPrep extract was incubated for 10 min at 95 °C, and thereafter processed according to the FASP protocol (in Microcon YM-30). Tryptic peptides were desalted using STAGE spin tips, and redissolved in 30 μl loading solution consisting of 2 % (v/v) acetonitrile and 0.05 % (w/v) trifluoroacetic acid in water. Approximate peptide concentrations were determined using a NanoDrop 2000 instrument (ThermoFisher Scientific, MA) and the A205 protocol (88).

### LC-MS/MS analysis

Approximately 1 μg of tryptic peptides was loaded onto a trap column (Acclaim PepMap100, C18, 5 µm, 100 Å, 300 µm i.d. × 5 mm) and then backflushed onto a 50 cm × 75 μm analytical column (Acclaim PepMap RSLC C18, 2 μm, 100 Å, 75 μm i.d. × 50 cm, nanoViper) for liquid chromatography-mass spectrometry (LC-MS/MS) analysis. A 120 minutes gradient from 4 to 40 % solution B (80 % [v/v] acetonitrile, 0.1% [v/v] formic acid) was used for separation of the peptides, at a flow rate of 300 nl/min. The Q-Exactive mass spectrometer was set up as follows (Top12 method): a full scan (300-1600 m/z) at R=70.000 was followed by (up to) 12 MS2 scans at R=35000, using an NCE setting of 28 eV. Singly charged precursors and precursors with charge>5 were excluded for MS/MS. Dynamic exclusion was set to 20 seconds.

### Bioinformatics

The raw MS data were analyzed using MaxQuant (89) version 1.5.3.40 and proteins were identified and quantified using the MaxLFQ algorithm (90). The data were searched against the UniProt *R. toruleoides* proteome (UP000016926; 8,138 sequences) supplemented with common contaminants such as human keratin and bovine serum albumin. In addition, reversed sequences of all protein entries were concatenated to the database to allow for estimation of false discovery rates. The tolerance levels used for matching to the database were 4.5 ppm for MS and 20 ppm for MS/MS. Trypsin/P was used as digestion enzyme and 2 miscleavages were allowed. Carbamidomethylation of cysteine was set as fixed modification and protein N-terminal acetylation, oxidation of methionines and deamidation of asparagines and glutamines were allowed as variable modifications. All identifications were filtered in order to achieve a protein false discovery rate (FDR) of 1%. Perseus version 1.5.2.6 (91) were used for data analysis, and the quantitative values were log_2_-transformed, and grouped according to carbon source. Proteins were only considered detected if they were present in at least two replicates on one of the two carbon sources. Missing values were imputed by drawing numbers from a normal distribution centred around the level of detection of the mass spectrometer. Student’s t-tests including permutation-based FDR (p<0.05) were used to detect significant differences between groups and volcano plots were constructed by plotting the negative logarithm of the p-values against the fold difference between two groups in a scatter plot, as shown in Additional file 1: Fig. S1. Differential expression between groups was assigned if proteins had a significant t-test and showed ≥ 2-fold difference. Due to increased variation in growth dynamics at early stationary phase, we used the p values (p≤0.05) instead of permutation-based FDR for t-test truncation to detect differences in cells cultivated with glucose or xylose as sole carbon source at the late stage of cultivation. Individual proteins were referenced by their UniProt accession numbers as described for the *R. toruloides* NP11 genome annotation (25). Gene set analysis was performed as described earlier (60) using available functional genome annotation (25).

## Declarations

### Authors’ contributions

IAT provided major contributions to study design and performed a major part of laboratory work, data analysis and writing the manuscript. JBr, JB and SS performed chemical analysis, data evaluation and was involved in manuscript writing. NM performed parts of the laboratory work and data analysis. MSk and MOA performed major parts of laboratory work. JN provided a major contribution for study design, data analysis and final manuscript writing. MS and EK coordinated the project and provided major contributions to study design, data analysis and final manuscript writing. All authors reviewed the final manuscript.

## Acknowledgements

We thank Dr. Tomas Linder for proofreading the manuscript.

## Availability of data and materials

All the data analyzed in this study are included in this manuscript and its additional files. The proteomics data has been deposited to the ProteomeXchange consortium (http://proteomecentral.proteomexchange.org) via the PRIDE partner repository (92) with the dataset identifier PXD012332.

## Consent for publication

Not applicable.

## Ethics approval and consent to participate

Not applicable.

## Additional files

**Additional file 1: Fig. 1.** GO term enrichment analysis and volcano plots of differentially expressed proteins in *R. toruloides* during either exponential growth phase or lipid production phase in medium containing either glucose (A) or xylose (B). For each GO term showing significant change (rank score of ≥ 5), the direction and significance of the relative changes in protein levels are shown, together with the total number of proteins within each GO term.

**Additional file 1: Fig. 2.** Volcano plots of differentially expressed proteins in *R. toruloides* cultivated on either glucose or xylose during early exponential growth phase (A), late exponential growth phase (B), or lipid accumulation phase (C)

**Additional file 2.** Complete list of proteins detected during batch cultivation of *R. toruloides* in minimal medium containing either glucose or xylose and their relative expression values

**Additional file 3.** Relative levels of differentially expressed proteins in glucose-grown *R. toruloides* during either exponential cultivated phase or lipid production phase.

**Additional file 4.** Relative levels of differentially expressed proteins in xylose-grown *R. toruloides* during either exponential cultivated phase or lipid production phase.

**Additional file 5.** Relative levels of differentially expressed proteins in *R. toruloides* cultivated on either glucose or xylose during early exponential growth phase

**Additional file 6.** Relative levels of differentially expressed proteins in *R. toruloides* cultivated on either glucose or xylose during late exponential growth phase

**Additional file 7.** Relative levels of differentially expressed proteins in *R. toruloides* grown on either glucose or xylose during lipid accumulation phase

**Additional file 8.** Relative levels of proteins involved in central carbon metabolism of *R. toruloides*.

