## Supplemental File 1 for "Proteome analysis of xylose metabolism in *Rhodotorula toruloides* during lipid production"

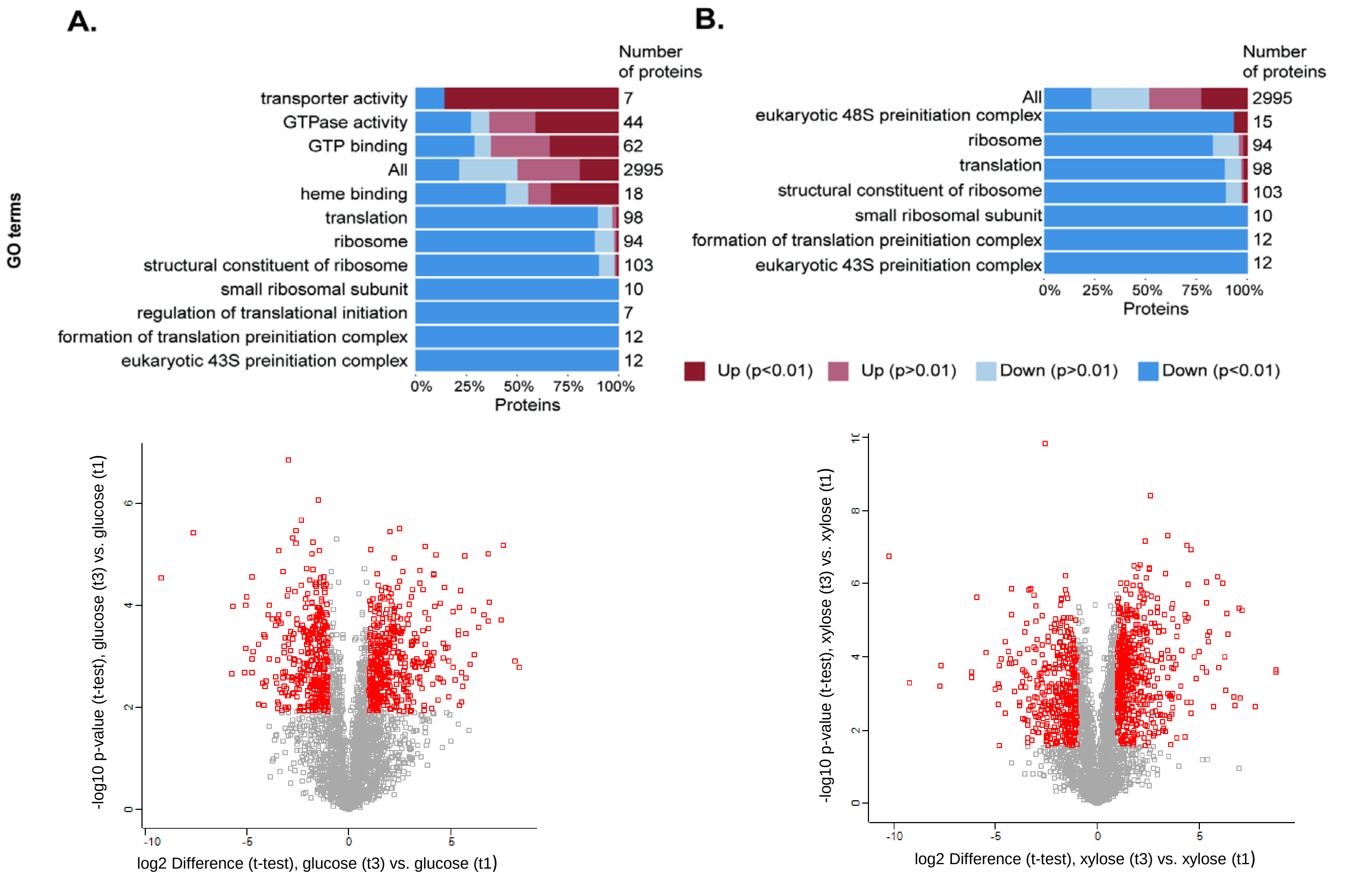


**Fig. S1.** GO term enrichment analysis and volcano plots of differentially expressed proteins in *R. toruloides* during either exponential growth phase or lipid production phase in medium containing either glucose (A) or xylose (B). For each GO term showing significant change (rank score of ≥ 5), the direction and significance of the relative changes in protein levels are shown, together with the total number of proteins within each GO term.

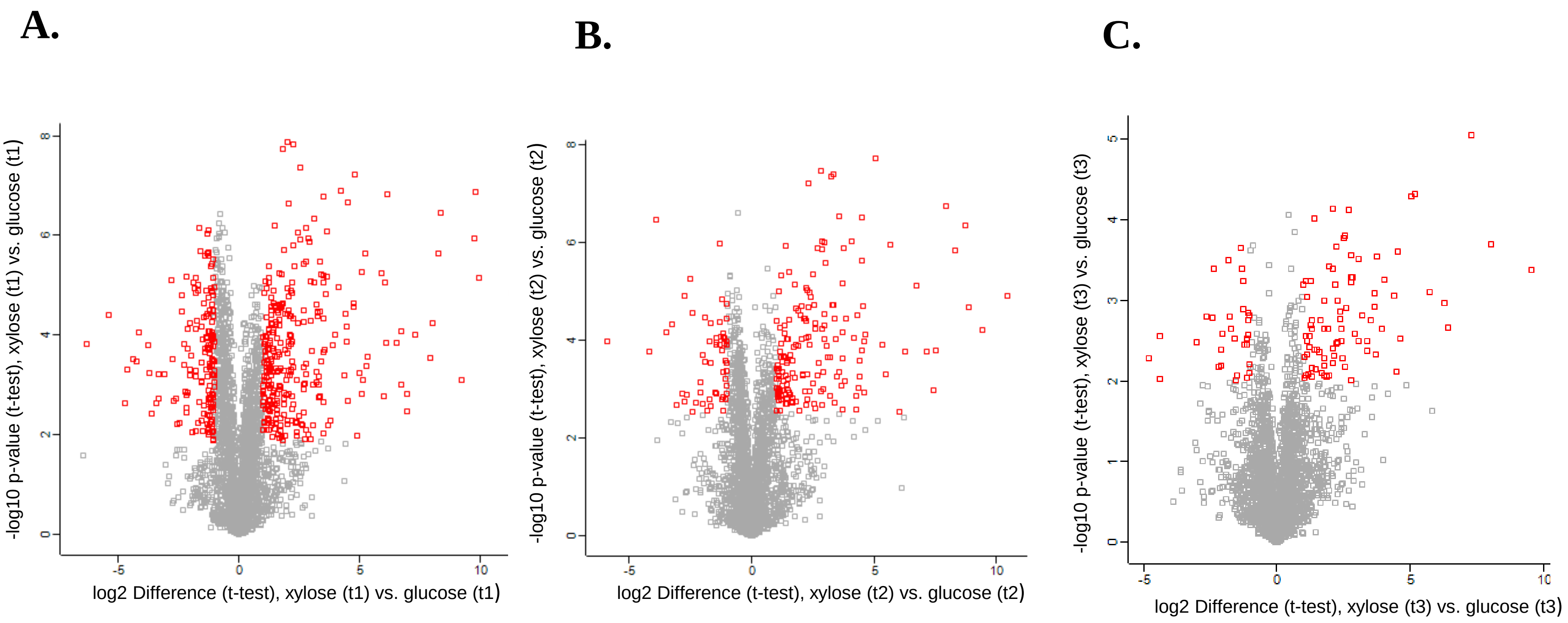


**Fig. S2.** Volcano plots of differentially expressed proteins in *R. toruloides* cultivated on either glucose or xylose during early exponential growth phase (A), late exponential growth phase (B), or lipid accumulation phase (C)
